# Gene transcription as a limiting factor in protein production and cell growth

**DOI:** 10.1101/626242

**Authors:** Eyal Metzl-Raz, Moshe Kafri, Gilad Yaakov, Naama Barkai

## Abstract

Growth rate and cell size are principle characteristics of proliferating cells, whose values depend on cellular biosynthetic processes in a way poorly understood. Protein production is critical for growth, and we therefore examined for processes limiting this production. Burdening cells with an excess of inert protein changed endogenous gene expression similarly to transcription-perturbing mutants, was epistatic to these mutants, but did not deplete respective factors from gene promoters. Mathematical modeling, corroborated by experiments, attributed this signature to a feedback which proportionally increases all endogenous gene expression, but lags at fast initiating genes already transcribed close to the maximal possible rate. As a possible benefit of maximizing transcription rates, we discuss a conflict between cell growth rate and size, which emerges above a critical cell size set by transcript abundance. We propose that biochemical limits on protein and mRNA production define the characteristic values of cell size and division time.

## Introduction

Cells tightly control the rate by which different proteins are produced. Regulation of protein expression can be direct, by factors that bind specific regulatory regions, or indirect, through changes in global parameters such as growth rate or cell size that subsequently alter protein expression. Indirect regulation may also occur through the depletion of common resources: when a general machinery is limiting, inducing the expression of specific proteins reduces production of other proteins that compete for the same machinery. Direct regulation of gene expression is extensively studied. By contrast, little is known about indirect regulations within the transcription and translation networks.

Ribosomes present a potentially limiting resource that determines the rate of protein translation and cell growth. Theory shows that in order to maximize growth rate, cells must tune their proteome compositions, so that they express the maximal number of ribosomes which can still be simultaneously engaged in translation. Under these conditions, in which all expressed ribosomes are constantly translating, growth rate is proportional to ribosome concentration, namely the ribosome:protein ratio. This predicted linear relation between ribosome concentration and cell growth rate was verified experimentally in a large number of cell types and growth conditions(Bremer and Dennis, 1996; Cox, 2003; Dennis et al., 2004; Emilsson and Kurland, 1990; Klumpp et al., 2013; Maaloe and Kjeldgaard, 1966; Marr, 1991; Metzl-Raz et al., 2017; Schaechter et al., 1958; Scott et al., 2010; Stoebel et al., 2008; Vind et al., 1993; Zaslaver et al., 2009). Similar considerations defined an upper limit on growth rate, which is obtained when all ribosomes are engaged in the translation of other ribosomes. Clearly, this theoretical maximum cannot be realized in cells, since additional proteins other than ribosomes are required for cellular activities. Remarkably, though, measured growth rate in rapidly growing yeast and bacteria fall in the range (30-50%) of this predicted maximum (Metzl-Raz et al., 2017; Scott et al., 2010).

Cells therefore function close to the predicted optimal growth, at which all expressed ribosomes are constantly engaged in translation. Still, recent analyses revealed important deviations from this predicted behavior, showing that cells maintain a fraction of non-translating ribosomes, likely as a reserve employed when translation demands unexpectedly increase (Korem Kohanim et al., 2018; Li et al., 2018; Metzl-Raz et al., 2017; Mori et al., 2017). Processes other than translation could therefore be limiting for protein production and cell growth. In support of this possibility, analyzing the growth phenotype of cells engineered to express excessive amounts of proteins demonstrated that both protein translation and gene transcription can become limiting, depending on growth conditions (Kafri et al., 2016).

The slow growth rate of cells forced to express excess proteins indicates that these cells experience some internal limitations. The molecular or mechanistic basis of these limitations, however, remained unclear. Gene expression signature is a sensitive probe of internal limitations; Budding yeast experiencing amino acids limitation, for example, induce GCN4-target genes (Gasch et al., 2000; Tamari et al., 2014), while cells depleted of internal phosphate induce PHO4 targets (Gurvich et al., 2017; Gutteridge et al., 2010). Both responses are gradual, providing a quantitative readout of the limitation. More generally, gene expression profiles of thousands of viable deletion mutants are available (Kemmeren et al., 2014; O’Duibhir et al., 2014) providing a readout for the internal consequences of perturbing a wide spectrum of cellular processes.

Motivated by this, we examined the gene expression changes induced by budding yeast cells forced to produce high amounts of mCherry protein, approaching ≈30% of the proteome. The transcription signature inflicted by this forced expression had little correlation with that of cells deleted of translation factors, including ribosome components. Rather, correlation was highest with mutants of the general transcription machinery, including the Mediator (specifically the Head and Tail sub-complexes), the SAGA, and the SWI/SNF complexes. We confirmed the phenotypic relevance of this correlation by demonstrating epistatic interactions between the burden and mediator mutants. Of note, only ∼5% of the DNA-bound mediator localized to the burden integration locus, refuting the possibility that the burden phenotype resulted from competition for limiting mediator.

To interpret our results, we considered a model of gene transcription. Since the mediator plays a major role in transcription initiation and re-initiation, we focused on the rate by which transcription is initiated. We discuss why transcription cannot be initiated at rates that exceed an upper bound, and review data reporting genes transcribed at rates close to this limit. We provide evidence that the transcription profile of the burden cells is attributed to these rapidly transcribed genes: burden initiates a feedback in cells, which proportionally increases the amounts of endogenous mRNAs, but fails to do so at genes already transcribed at high rates that approach the maximal limit.

Our results therefore suggest that cells drive transcription rates of rapidly transcribed genes to their maximal possible rate, and this is accentuated in the burden cells. But what could be the benefit of working at this limit? As a possible answer, we consider the ribosome-centered model of cell growth, in which growth rate is proportional to the concentration of translating ribosomes and cell size is proportional to the total protein amount. Within this model, we describe a critical cell size, above which any further increase in size compromises growth rate. This critical size depends on the number of transcripts, which in turn, defines the maximal number of bound ribosomes that can be translating simultaneously. Maximizing transcription rates therefore allows increasing cell size without compensating cell growth. Based on these results, we propose that biochemical limits on protein and mRNA production set the characteristic values of cell size and division time.

## Results

### The transcriptional response to protein burden

Cells modify their gene expression when subject to nutrient limitations or when key processes are perturbed. The expression signature of cells forced to express excessive amounts of inert protein could therefore report on the internal limitations inflicted by this burden. We previously constructed a library of budding yeast cells expressing an increasing copy number (1-20) of genomically integrated pTDH3-mCherry constructs. These strains produce mCherry proteins at increasingly high levels (Figure 1A), approaching ≈30% of the total cellular proteins (Kafri et al., 2016). To define the transcription changes inflicted by the burden, we grew cells from this library to logarithmic phase and measured their gene expression. We repeated this profiling experiment in three conditions: standard media (SC), media low in nitrogen and media low in phosphate. As expected, the overall pattern of gene expression changed gradually with mCherry amounts (Figure 1B-C).

**Figure 1.**
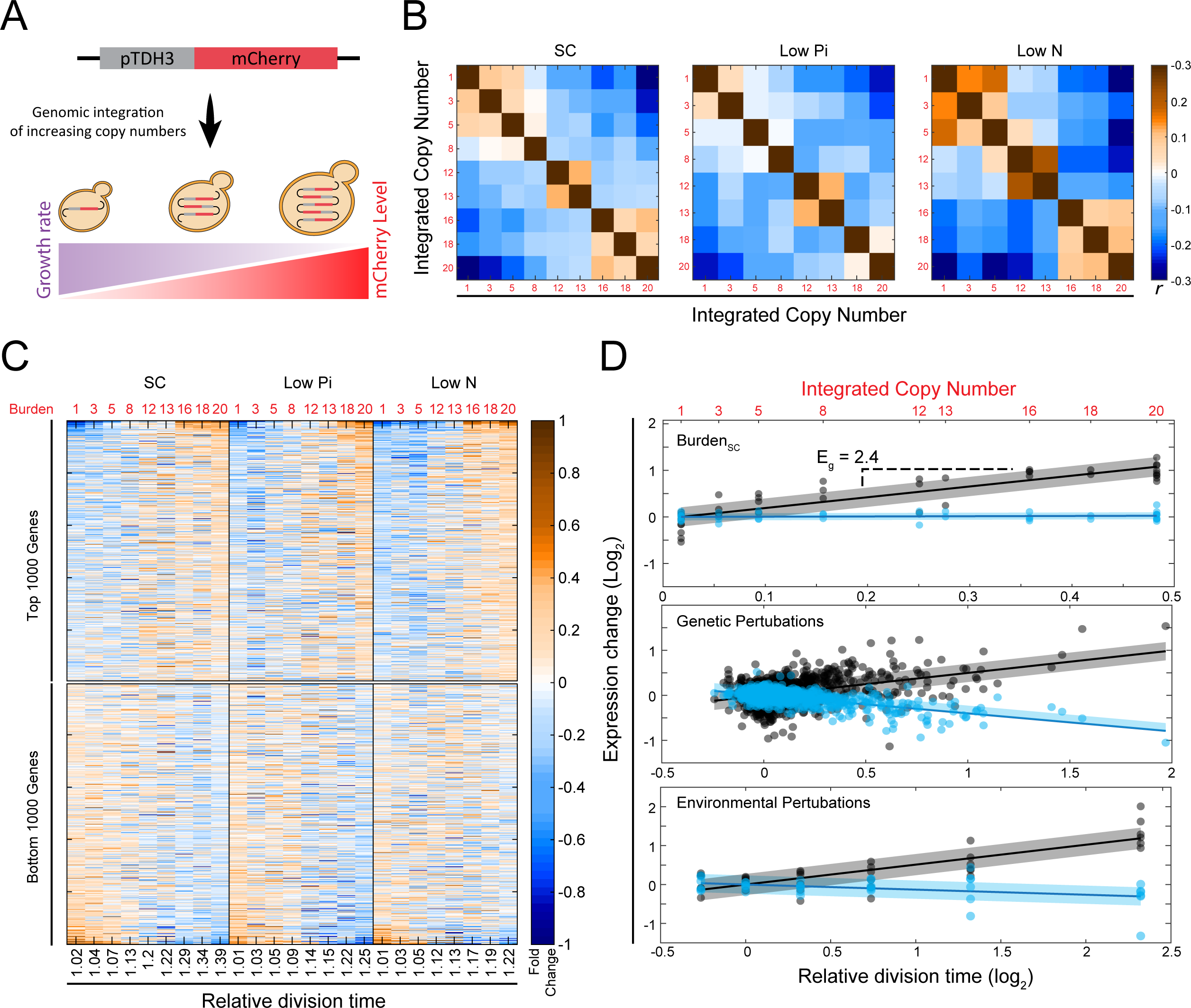
The transcription signature of burden cells: (A) *Engineering libraries of burdened cells*: Schematic - Transforming yeast with a linearized plasmid harboring *TDH3*-promoter driven mCherry results in a variable number of genomic tandem integrations per cell. Individual clones with increasing copy number are taken for further analyses. See methods for details. (B) *Cellular transcription response changes gradually with the level of forced expression:* Shown are the Pearson correlations *r* between the transcription profiles of cells burdened with the indicated mCherry copy number, grown in the indicated conditions. Expression levels were normalized by their mean value in the specific condition and log_2_-transformed. (C) *Expression levels*: The top and bottom 1000 affected genes were selected and sorted by the strongest change with the relative growth rate in standard media SC. Expression is shown as a function of cell growth rate (x-axis) relative to WT. (D) *Growth-rate response (“E*_*g*_*”):* Shown are the expression levels of SIS1 (black) and HHT2 (blue) measured in strains of the three indicated datasets as a function of the strain’s growth rate. The gene-specific growth-rate responses are defined by the slope of this relation, as indicated.

### Distinguishing expression changes specific to protein burden from changes common to slow-growing cells

Previous studies described genes whose expression correlates with growth rate over a wide range of genetic or environmental perturbations (Brauer et al., 2008; Gasch et al., 2000; Hughes et al., 2000; Levy et al., 2007; O’Duibhir et al., 2014; Regenberg et al., 2006; Zurita-Martinez and Cardenas, 2005). Since forced expression of unneeded proteins reduces growth rate in proportion to the added burden (Dong et al., 1995; Kafri et al., 2016; Scott et al., 2010; Shachrai et al., 2010), transcription changes observed in these cells could result from their slow-growth phenotype. To distinguish expression changes that are specific to the burden from those that are general consequences of slow growth, we compared our data to two published compendiums reporting transcription profiles and growth rates. The first dataset described wild-type cells grown in chemostat-based environments (“Environmental Perturbations” (Brauer et al., 2008)). The second described 1,484 viable deletion mutants grown in non-stress conditions (“Genetic Perturbations” (Kemmeren et al., 2014; O’Duibhir et al., 2014)). In each dataset, we defined the rate at which gene expression changes with growth rate (*E*_*g*_, Figure 1D, Figure S1A). This provided us with three directly comparable gene-specific measures for each dataset (“Burden”, “Genetic” and “Environmental” perturbations).

The growth-rate responses observed in the two external genetic and environmental perturbation datasets were highly correlated (Figure 2A). By contrast, the burden response was notably different (Figure 2B-C). The majority of expression changes observed in the burden cells therefore result specifically from the forced production of proteins. To understand these changes, we tested for classes of genes that are preferentially affected. For this, we checked enrichment of gene groups defined by GO-slim, binding to the same transcription factors (TFs) and co-expression in multiple datasets (Ihmels et al., 2002, 2004) (Figure 2D, Figure S2A). Hsf1-dependent chaperones were consistently induced in the burdened cells, likely indicating protein folding stress. Those genes, however, were also induced more generally in slow-growing cells. Apart from these, we did not detect any other group using this enrichment test. In particular, neither Gcn4-dependent genes, reporting on amino-acid limitations, nor oxidative-phosphorylation genes, related to energy balance, showed a consistent change with increasing burden. Indeed, the specific rates of glucose uptake and ethanol production remained invariant to the burden, suggesting that central metabolic fluxes remain largely invariant to the protein burden (Figure S2B-C).

**Figure 2.**
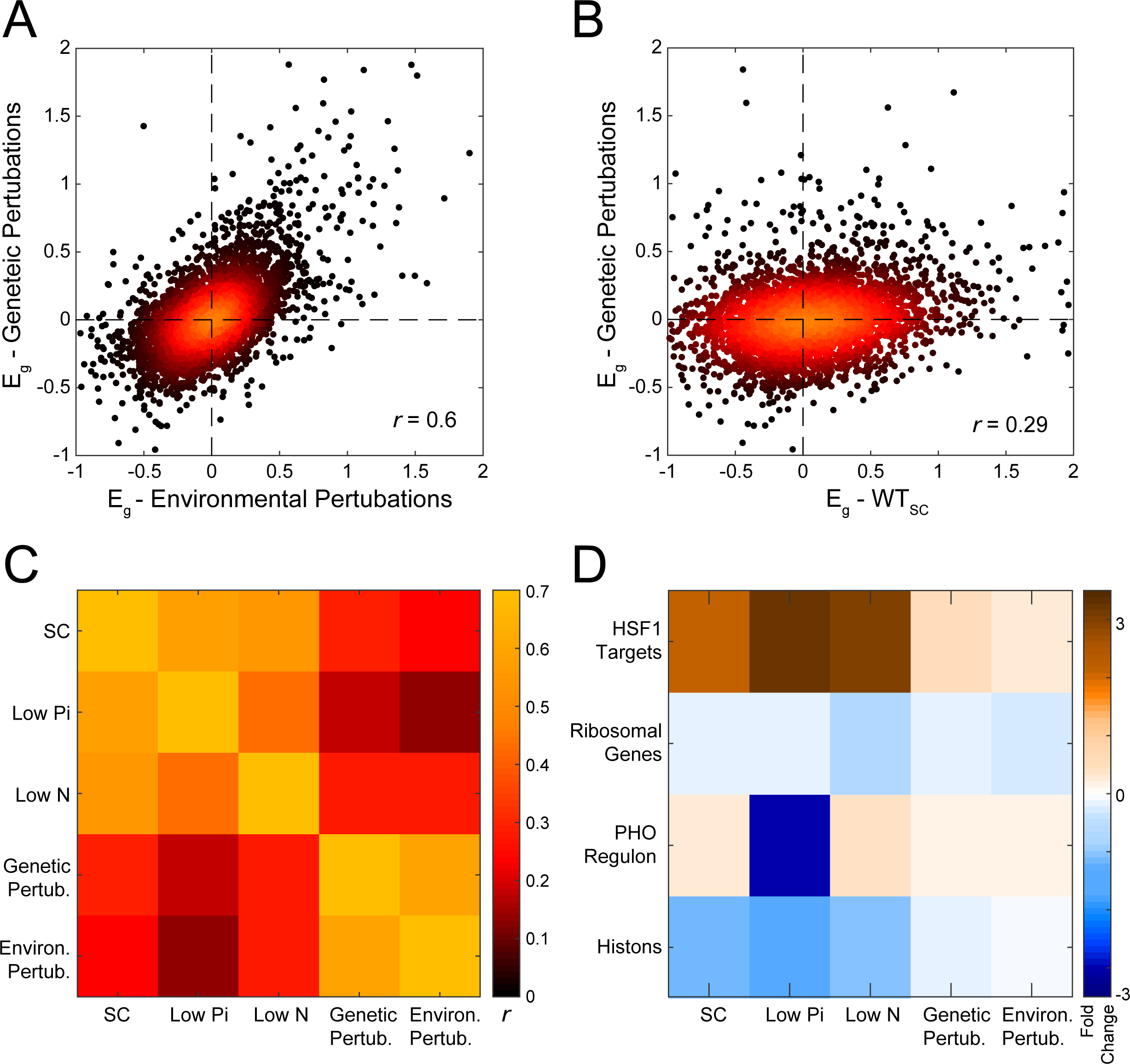
The transcription response to protein burden is distinct from the slow-growth program: (A-C) *Correlation between growth-rate responses of burden cells and perturbed cells:* Shown are the values of the indicated Growth-rate responses (A, B), and the respective Pearson correlations (C). (D) *Gene-groups showing a coherent growth-rate response:* The set of genes showing the most significant growth-rate response were defined for each dataset. These sets of genes were compared with pre-defined gene groups associated with a common function or regulatory properties. Shown are groups with significant enrichment in at least one gene-set (See also Figure S2A).

### Signatures of perturbed transcription initiation in burden cells

As a complementary approach to predict cellular processes perturbed in burden cells, we measured the correlations between the transcription changes caused by the burden and the transcription signatures of the 1,484 gene-deletion mutants(Kemmeren et al., 2014; O’Duibhir et al., 2014) (Figure 3A, Figure S3A). To control for growth-related changes, we also correlated the mutants with the growth-related transcription response (Figure 3A, “Growth Effect”). Correlations of burden response with translation-related mutants, including the deletion of ribosome components, were relatively low (Figure 3B, left; Figure S3B-C). Similar low correlations were seen with mutants associated with protein or mRNA degradation (Figure S3B).

**Figure 3.**
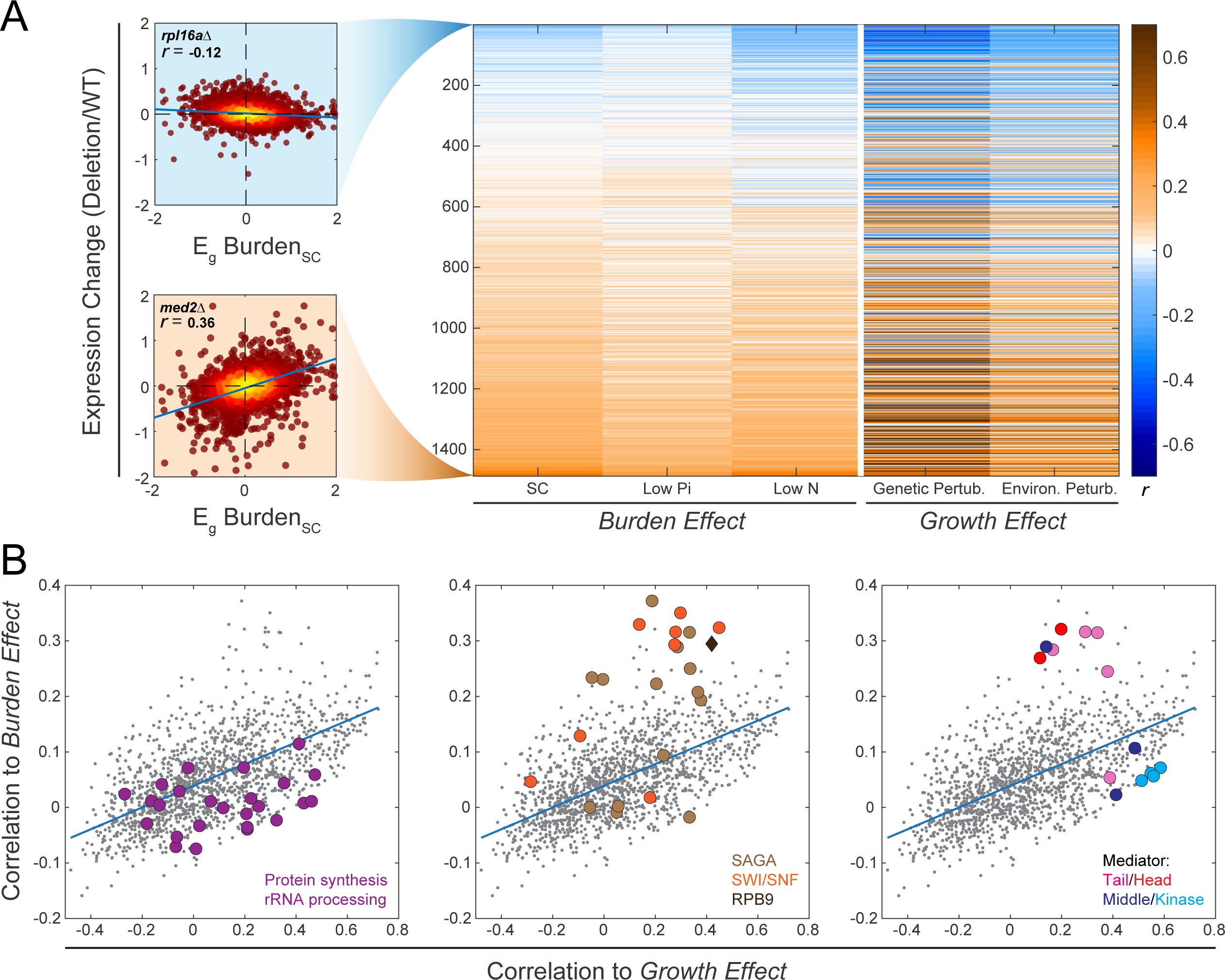
Transcription signature of burden cells correlates with that of transcription-perturbing mutants: (A) *Correlations between the burden response and the transcription response to gene-deletion mutants:* Shown are the Pearson *r* correlations between the growth-rate response E_g_ (measured in the indicated dataset and condition), with the transcription signature of each individual gene-deletion mutant. Mutants are ordered by the correlation values with the burden response, averaged over the three conditions. Specific mutants are highlighted, as indicated (see also Figure S3A). (B) *Distinguishing mutants that correlate specifically with the burden response:* Correlation between mutant signature and burden response (as in A, averaged over three conditions) are plotted as a function of the correlations between mutant signature and growth-rate response (as in A, averaged over the genetic and environmental responses). Each dot is a mutant, color-coded as indicated. See also Figure S3B.

The majority of mutants that correlated preferentially with the burden cells were involved in gene transcription. These include the deletion of RPB9, the only non-essential component of the RNA-Polymerase II profiled in the compendium, and most components of the chromatin-remodeling complexes SAGA and SWI-SNF. Particularly high correlations were found for mutants of the Mediator complex (Figure 3B, middle and right; Figure S3B,D). The Mediator plays a central role in transcription initiation by physically linking specific transcription factors (TFs) with the general machinery. It is composed of a tail sub-complex, which binds specific Upstream Activating Sequences (UASs) and recognizes specific transcription factors, a head sub-complex, which binds RNA polymerase II, and a middle sub-complex that bridges the head and tail sub-complexes(Allen and Taatjes, 2015; Ansari and Morse, 2013; Haberle and Stark, 2018; Jeronimo and Robert, 2017; Malik and Roeder, 2016; Sikorski and Buratowski, 2009; Soutourina, 2017; Tantale et al., 2016). An additional inhibitory sub-complex of the Mediator, the Kinase, has a mutually exclusive binding with the polymerase and dissociates upon polymerase binding and transcription initiation(Andrau et al., 2006; Aristizabal et al., 2013; Clark et al., 2015; Gonzalez et al., 2014; Jeronimo et al., 2016; Petrenko et al., 2016). Of the seven components of the Mediator tail or head sub-complexes profiled in the compendium, six were correlated with the burden response. By contrast, mutants of the middle or inhibitor sub-complexes showed no such correlation (with the exception of MED31 whose subunit association is somewhat unclear (van de Peppel et al., 2005)).

To examine whether these similarities in gene expression result from shared internal perturbations, we focused on the Mediator. First, we re-engineered the respective mutants and profiled their transcriptomes, verifying their correlations with the burden strains (Figure S3B-D). Second, we examined whether burden cells show phenotypes similar to those exhibited by mutants of the mediator head or tail sub-complexes. Cells deleted of the Mediator Kinase inhibitory sub-complex are pseudo-hyphal and flocculate when growing in liquid media (Hengartner et al., 1995; Holstege et al., 1998). This phenotype is reverted by deleting components of the Mediator tail or head sub-complexes (Gonzalez et al., 2014; Jeronimo et al., 2016; Law et al., 2015; Palecek et al., 2000; van de Peppel et al., 2005)(Figure 4A, Figure S4A). We therefore asked whether protein burden similarly reverts the flocculation phenotype of kinase-deleted cells. This was indeed the case: increasing mCherry expression in kinase*-*deleted cells progressively reduced flocculation (Figure 4A-B, Figure S4A). Therefore, protein burden phenocopies the mediator tail or head mutant phenotype, consistent with their similarity in gene expression.

**Figure 4.**
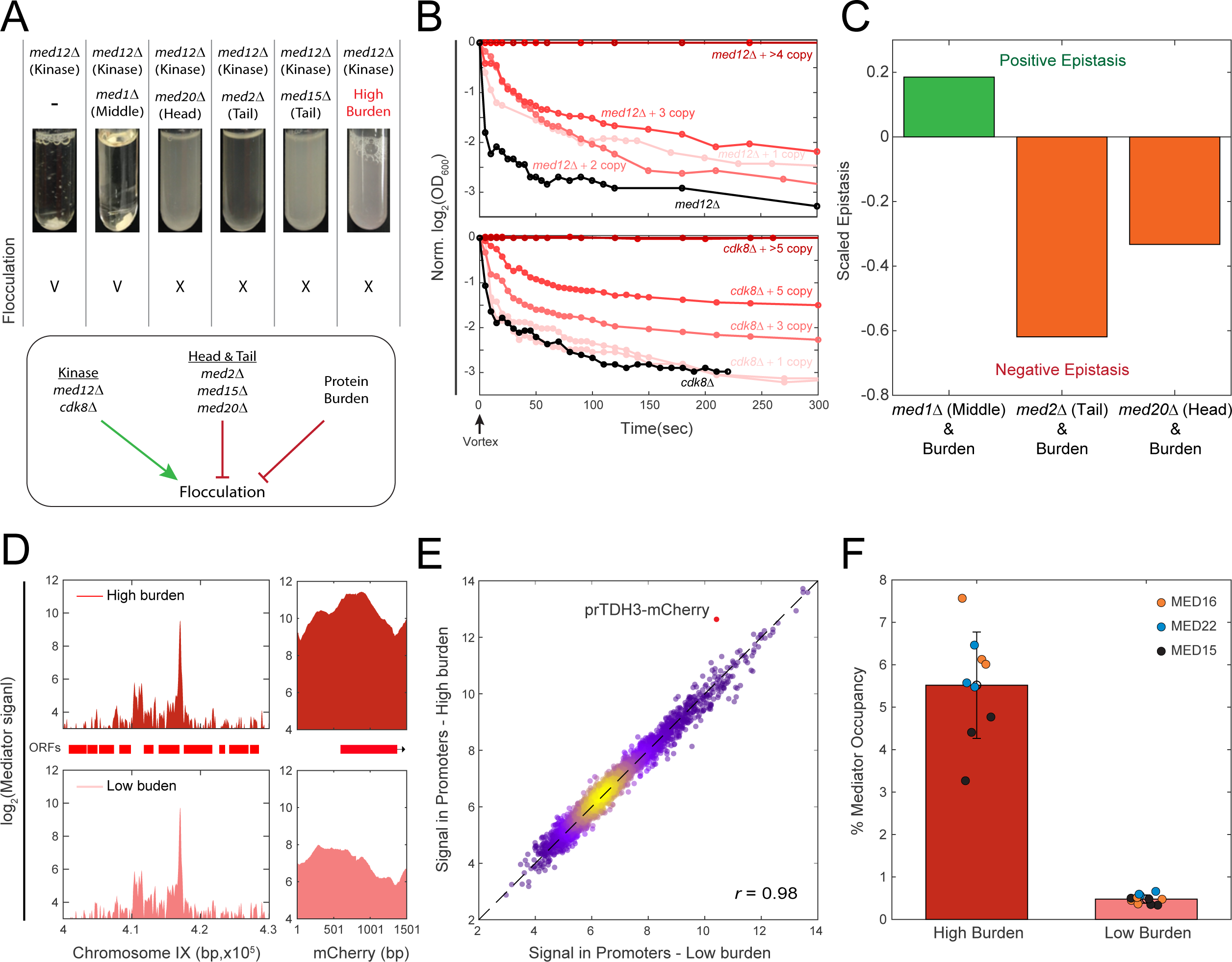
Genetic interaction between burden and Mediator mutants: (A-B) *Protein burden phenocopies mutants deleted of mediator head or tail components:* Shown are flocculation phenotypes, quantified as described in Methods. Deletion of the mediator kinase subunit *MED*12 induces flocculation, but this is reverted when deleting components of the head or tail sub-complexes, or by introducing protein burden. Note the gradual effect of increasing burden in this phenotype (B and Figure S4A). (C) *Epistatic interactions between burden and mediator mutants:* Burden libraries were prepared in the background of the indicated mutants, and growth rates were quantified using sensitive competition assays. Epistatic interactions were defined according to Segrè et al. (2005) (methods). (D-E) *Binding of mediator to endogenous promoters is invariant to protein burden:* Genomic binding profiles of the indicated mediator components were profiled in high and low-burden strains using ChIP-Seq. Read coverage along chromosome IX and at the mCherry ORF are shown in (D), and promoter-averaged binding strengths in the high vs. low burden strains are shown in F. Note the increased binding to the *TDH3*-mCherry promoter in the high-burden cells. The fraction of mediator that binds to the burden constructs is shown in F.

Mutants that affect the same process often exhibit epistatic interactions (Elena and Lenski, 1997; Hartman et al., 2001; Lenski et al., 1999; Phillips, 2008). To examine whether this is also the case for protein burden and Mediator mutants, we prepared burden libraries in the background of mediator mutants (Figure S4B-C,F). We measured the relative fitness of cells in these libraries, and quantified their epistatic interactions based on Segrè et al. (2005) (Methods). Negative epistasis was observed between the burden and mediator tail or head mutants, consistent with the similarity in their transcription profiles. As expected, this interaction was lost in a mutant of the middle sub-complex, which in fact showed a positive (alleviating) interaction with the burden (Figure 4C, Figure S4D). The pattern of epistatic interactions between burden and mediator mutants is therefore consistent with the similarities in their gene expression profiles.

### Mediator binding to endogenous promoters remains invariant in burden cells

The similarities in gene expression and phenotypes between burden cells and mediator mutants may be explained if mCherry production depletes the Mediator from endogenous promoters. To examine this, we measured the genome-wide binding profiles of three Mediator head and tail subunits using ChIP-Seq. Binding patterns at endogenous genes were insensitive to the burden (Pearson Correlation of 0.98 Figure 4D-E, Figure S4E). Further, even in the strains that expressed ≈15 copies of the mCherry gene and showed ≈25% growth defect, only ≈5.5% of detected binding events were localized to the integration locus of the mCherry-expressing construct (Figure 4F). Therefore, we find it unlikely that the burden transcription signature results from this modest depletion.

### Pol2 elongation rate defines a maximal biochemical limit for the rate of gene expression

To examine for a complementary explanation for the similarity between the burden and mediator phenotypes, that does not invoke depletion of mediator from endogenous promoters, we considered a model of gene expression. The Mediator promotes initiation and re-initiation (Andrau et al., 2006; Aristizabal et al., 2013; Haberle and Stark, 2018; Harlen and Churchman, 2017; Jeronimo et al., 2016; Soutourina, 2017), and we therefore examined the physical limits constraining initiation rates. It was previously noted that the rate of transcription initiation is limited by the time required for the polymerase to elongate away from its initiation site (Choubey et al., 2015; Ehrensberger et al., 2013) (“Pol2 Clearance”). Considering the polymerase footprint on DNA of ∼35bp (Brabant and Acheson, 1995; Selby et al., 1997), this imposes a maximal initiation rate of ∼1.2 seconds/transcript (corresponding to an average elongation rate of 2 kb/min (Darzacq et al., 2007; Edwards et al., 1991; Koš and Tollervey, 2010; Mason and Struhl, 2005; Pelechano et al., 2010; Pérez-Ortín et al., 2007; Swinburne and Silver, 2008; Zenklusen et al., 2008), Figure 5A, Figure 5A - Supplementary file 2). Initiation rates are expected to vary widely between genes, depending on their expression levels and burst frequencies, with measured initiation rates available for few genes only. Still, several of these measurements report initiation rates that are on par with this maximal limit: The *Drosophila* hsp70 transcript, for example, is produced every ∼1.5-3 seconds (Lengyel and Graham, 1984), similar to the production rate of the Dictostylium Act1 gene during transcription bursts(Corrigan et al., 2016). In budding yeast, oxidant-exposed cells produce TRR1 transcript at estimated four second intervals (Monje-Casas et al., 2004). Similar rates were measured for the PDR5 gene during its transcription bursts (Zenklusen et al., 2008), and estimated for HIS1 transcripts driven by strong promoters (Iyer and Struhl, 1996). Further, estimating initiation rates based on measured values of mRNA abundance and degradation rates are also consistent with these high initiation rates, suggesting that highly expressed genes, and in particular those which are produced in bursts, are transcribed at rates that approach the theoretical maximum (Figure 5A - Supplementary file 3).

**Figure 5.**
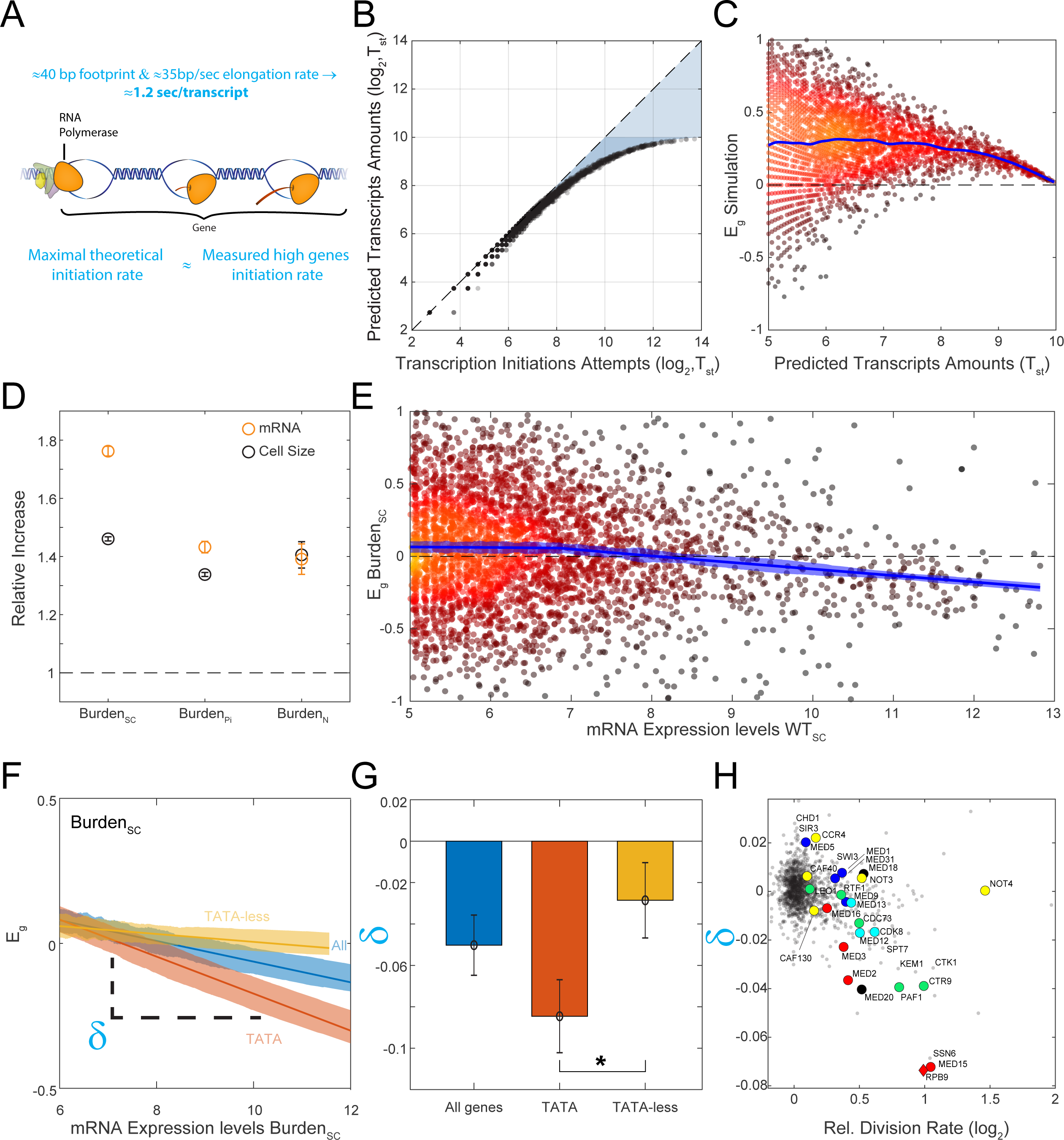
Transcription-promoting feedback activated in burden cells: (A) *The maximum possible limit of transcription initiation rate:* A new initiation event can only occur once the polymerase has elongated away from its initiation site. This elongation rate therefore defines an upper bound on the possible rates of transcription initiation. (B–C) *Simulation of the transcription initiation process:* The model assumes that initiation attempts are stochastic, characterized by some attempt rate. An attempt is deemed successful if it occurs at a sufficient delay from a previous successful attempt. This delay corresponds to the time required for the polymerase to clear the initiation site. Shown are the frequency of successful initiation events as a function of the attempt rate (B). The consequence of increasing the overall attempts frequency, as we assume happens in burden cells, is shown in (C), where the blue line is cubic smoothing spline. Note the limited efficiency of this feedback at genes transcribed at high rates. (D) *Burden cells increase the overall amounts of endogenous transcripts:* Total amount of mRNA was measured using sequencing, calibrated by an external spike-in reference. (E-G) *Relative expression of high-abundance genes tends to decrease in burden cells:* Shown are the relative changes in gene expression in burden cells as a function of absolute mRNA abundance in wild-type cells (E(. This effect is accentuated in TATA containing genes (F), with the lines showing the linear fit and the shading the SE. The expression-dependent bias was quantified by the slopes (δ) of these dependencies. Averaged values of this bias, calculated in the different datasets are also shown (G). Error bars represent SE. *Mutant strains preferentially affecting highly-expressed, TATA-containing genes:* The expression-dependent bias (δ) was calculated for each deletion mutant, as in E-F. Shown are the values of this bias, as a function of the mutant growth rate. Genes associated with transcription initiation and elongation are marked.

Therefore, the rate of transcription elongation seems to limit the rate of gene expression (Choubey et al., 2015; Ehrensberger et al., 2013). The existence of such a limit implies that not every attempt to initiate transcription is successful, even at attempt frequencies below the theoretical limit. This is captured by our simulations, where we assumed varying rates of initiation attempts, and measured the frequency of successful events. Indeed, the rate of successful initiation events approaches saturation at frequencies significantly lower than the theoretical maximal rate (Figure 5B).

### Endogenous transcript levels increase proportionally in burden cells, but less so in highly expressed, TATA containing genes

Is this limit on transcription initiation, described above, relevant for explaining the common transcription changes observed in the mediator and burden mutants? We reasoned that this may be the case if, in an attempt to retrieve homeostasis, burden cells would activate a global feedback to increase their overall transcription capacity. Such a feedback, corresponding to increased rates of initiation attempts at all genes, will lead to a proportional increase in the expression of most genes, but will be less effective in genes already transcribed close to their maximal limit (Figure 5C).

To examine if burden cells increase their overall expression of endogenous genes, we compared the total mRNA amounts using an *S. paradoxus* spike-in as a normalization standard. As we hypothesized, mRNA content in the burden cells was significantly (≈75%, SC) higher than in wild-type (Figure 5D). Therefore, the majority of genes show a proportional increase in abundance in the burden cells, to an extent that in fact exceeds the size increase of these cells.

Next, we wished to examine whether this increase in mRNA abundance of burdened cells is less pronounced in rapidly transcribed genes. We reasoned that rapidly initiating genes are enriched amongst highly expressed genes, and in particular among these genes that also contain a TATA-box which is often transcribed through bursts of rapid initiating events (Cho et al., 2016; Corrigan et al., 2016; Larsson et al., 2019; Zenklusen et al., 2008). Burden cells indeed preferentially reduced expression of highly expressed genes (Figure 5E), and this effect was accentuated in TATA-containing genes (Figure 5F-G, Figure S5A-B).

Consistent with the overall similarity in expression profiles, Mediator mutants deleted of the head or tail subunits showed a similar preferential effect on highly-expressed, TATA-containing genes (Figure 5H, Figure S5C-D). We attribute this preference to the mediator role in promoting initiation and re-initiation, which are increasingly important in rapidly initiating genes. Further, supportive of our hypothesis that this signature is associated with the maximal transcription initiation rate, the same signature was also found in slow-elongating polymerase mutants predicted to reduce the initiation rate and accentuate this limit (e.g. mutants of the PAF1 transcription elongation complex (Figure 5H, Figure S5C)). Finally, we verified that this signature was not a general consequence of the slow growth of these mutants (Figure S5A).

### The ribosome-centered model of gene expression predicts a critical cell size defined by transcript abundance, above which size begins to conflict with growth-rate optimality

Our results attribute the common phenotype of burden and mediator mutants to genes transcribed at rates that approach the maximal possible expression limit. The fact that cells maximize gene transcription rates, driving rapidly transcribed genes to rates that are close to the maximal limit, was initially surprising to us, since it is commonly assumed that mRNA abundance is irrelevant for cell growth. In the ribosome-centered model, for example, cells grow by means of ribosome-dependent protein translation (Bremer and Dennis, 1996; Dennis et al., 2004; Emilsson and Kurland, 1990; Klumpp et al., 2013; Maaloe and Kjeldgaard, 1966; Marr, 1991; Metzl-Raz et al., 2017; Schaechter et al., 1958; Scott et al., 2010; Vind et al., 1993). This leads to the experimentally observed exponential increase in protein content P, with specific growth rate that is proportional to the concentration *r* of translating ribosomes (R): r=R/P. Cell size is not considered explicitly. Rather, it is assumed that size increases in proportion to protein content P, a requirement which is indeed essential for maintaining growth homeostasis (Jorgensen et al., 2004; Levy et al., 2007; Lin and Amir, 2018; Moss, 2004). As mentioned in the introduction, maximizing growth rate requires that all expressed ribosomes are constantly translating. This also ensures that cells maintain exponential growth, since the fraction of translating ribosomes, and therefore growth rate, does not change with increasing size. Therefore, within this class of optimal-growth models, cell growth rate and size are completely independent.

We noted, however, that although mRNA abundance is not included explicitly in this model, it is in fact relevant to its dynamics since the abundance of the mRNA substrate constrains the maximal number of ribosomes, R^max^, that can be simultaneously bound and engaged in translation. By this, the abundance of transcripts also sets a critical cell size above which size increase begins to conflict with growth rate optimality. To see this, consider growing cells for which the optimal concentration of actively translating ribosome is *r=R/P.* This optimality can be maintained over a range of total protein content (cell size), provided that P<P^max^, where P^max^ =rR^max^: For cell size (∼P) smaller than this critical value, cells can maintain the same active concentration *r* of constantly translating ribosomes. By contrast, maintaining this optimality for larger cells requires that the number of co-translating ribosomes exceed the maximal number R^max^ allowed by the available mRNA (Figure 6).

**Figure 6.**
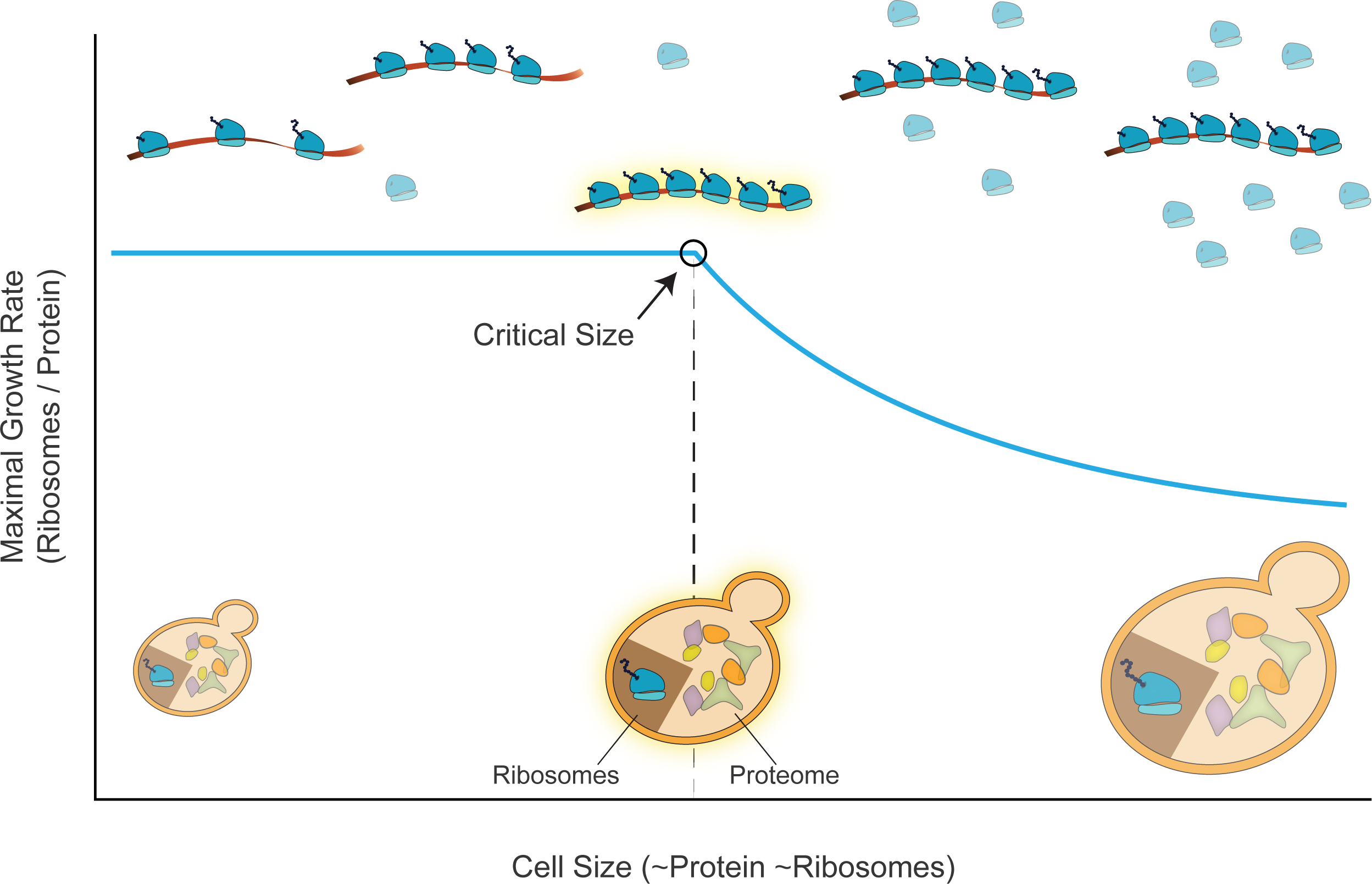
Proposed model for a critical cell size that depends on transcript abundance. We suggest a critical cell size above which increasing size (X axis, directly proportional to protein and ribosome abundance) begins to compete with optimal growth rate (Y Axis). We assume a fixed mRNA abundance and examine how changing the abundance of protein (equivalent of cell size) is expected to impact optimal growth. To maintain optimal growth, the ribosomal fraction scales with the abundance of proteins. As long as it is low enough, the fraction of translating ribosomes can also be maintained, as sufficient mRNA is available. In this regime, growth rate is not affected by the change in total protein levels. However, further increasing protein abundance beyond this size, necessarily reduces the fraction of co-translating ribosomes, leading to a reduction in cell growth rate. See text for details.

Maximizing the rate of mRNA production is therefore beneficial for increasing this critical size, thereby allowing cells to maintain an optimal growth rate over a broader range of sizes.

## Discussion

In this work, we set out to determine factors and processes that are limiting for protein production and cell growth. We approached this by comparing the transcription profiles of cells forced to express excessive levels of inert mCherry proteins with the respective signatures of hundreds of gene-deletion mutants. We initially expected that the need to translate high levels of mCherry proteins will deplete ribosomes from endogenous transcripts and will therefore imitate conditions of insufficient translation, corresponding perhaps to deletions of ribosome subunits. This, however, was not the case. Rather than translation-perturbing mutants, we found that the burden cells mostly resemble components of the general transcription machinery. We examined if this shared signature results from the depletion of the identified transcription components (e.g. Mediator complex) from certain endogenous promoters. This, however, was not the case, as only ∼5% of the bound mediator localized to the burden constructs.

Modeling the transcription process highlighted a limitation of a very different nature: a physical limit that restricts the maximal possible rate of transcription initiation. This limit depends on molecular properties of the polymerase: its DNA footprint and the rate by which it elongates along the transcript to clear the promoter for another incoming polymerase. Available data suggests that this limit is relevant for *in-vivo* transcription rates, as highly transcribed genes appear to be transcribed close to this limit. We found that protein-burden cells increase the amount of endogenous mRNA, perhaps as a way to retrieve wild-type protein concentration, but fail to do so at highly expressed, TATA-containing genes. Their transcription signature can therefore be explained by the inability to further induce the expression of genes that are already transcribed at high rates that are close to the maximal possible limit.

Why would cells transcribe genes close to this boundary of maximal transcription? Could there be a functional benefit in maximizing mRNA production? Following the accepted paradigm for modeling cell growth, we revealed a conflict between growth rate and cell size that emerges above a critical cell size: below this critical size, growth rate is independent of size, but above this critical size, further increasing size compromises optimal growth (Figure 6). Of note, in this analysis we assumed that cell size remains proportional to protein abundance, as is expected under conditions of homeostasis. This critical size then depends on the number of expressed transcripts, which defines the maximal number of ribosomes that can simultaneously engage in translation. Maximizing transcription rates may therefore allow more ribosomes to be expressed, thereby increasing cell size without compensating cell growth optimality.

Whether cells work close to this limit of maximizing the ribosome number is not clear: Budding yeast express an estimated 200,000 ribosomes, compared to 35,000 transcripts (supplementary text and (Miura et al., 2008)). If all mRNAs were to bind ribosomes at the same efficiency, this would amount to an average of ∼8 ribosomes per mRNA (Arava et al., 2003; Zenklusen et al., 2008). Considering the footprint of a ribosome on mRNA (∼35bp (Brabant and Acheson, 1995; Selby et al., 1997)), we expect a rather low density of ribosomes on most transcripts. The extent to which mRNA restricts ribosome number, however, should be evaluated based on the highest ribosome densities found at rapidly translated genes. Ribosome densities are higher at gene beginnings where elongation is slower. Indeed, it was estimated that 20% of ribosomes are positioned adjacent to another ribosome, being detected as a single footprint in ribosome profiling experiments (Diament et al., 2018). This high density, at least in some transcripts, may argue that ribosome number is adjusted to mRNA abundance, to fully utilize the available transcripts and maximize cell size.

Taken together, we propose that maximizing transcript production may serve to increase the maximal cell size (or protein content), for which cells can still maintain optimal growth. The maximal possible initiation rate, which limits this production, may therefore serve as a fundamental physical constraint, limiting cell size. This is analogous to the time of ribosome translation, which is the fundamental unit defining cell growth rate. These two physical constraints on transcription and translation, set by the basic biochemical parameters inherent to these processes, may therefore define the characteristic values of the division time and size of rapidly proliferating cells.

**Figure S1.**
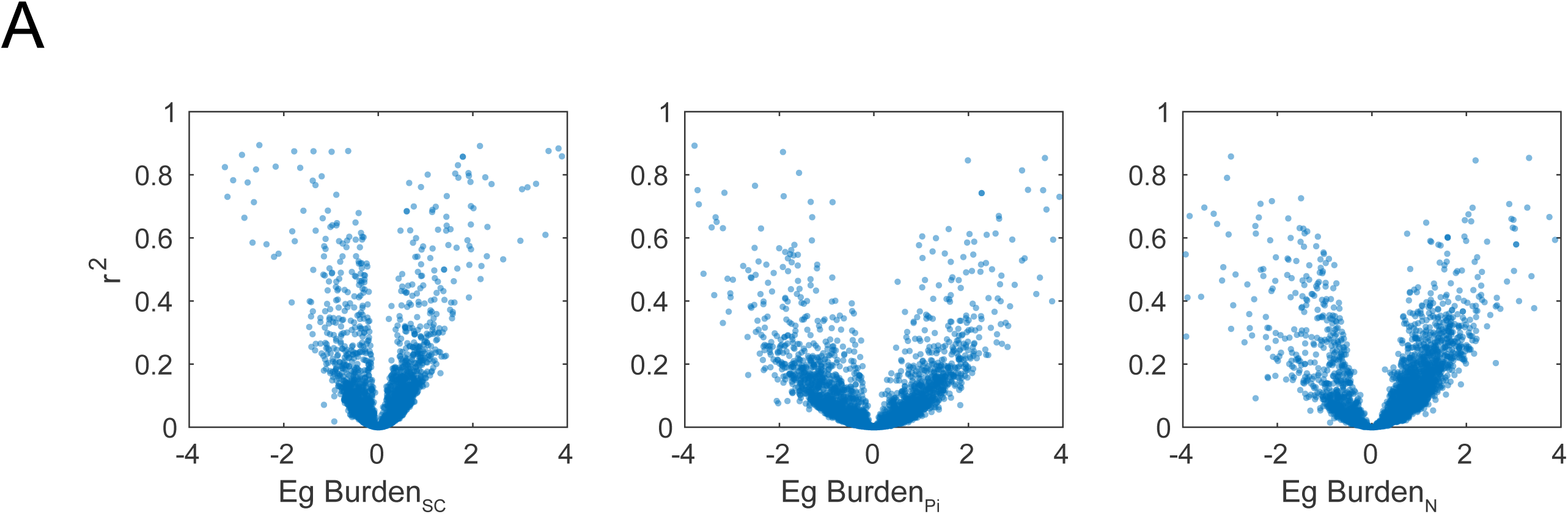
A Higher E_g_ calculated for burden in each of the three conditions corresponds to a higher r^2^.

**Figure S2.**
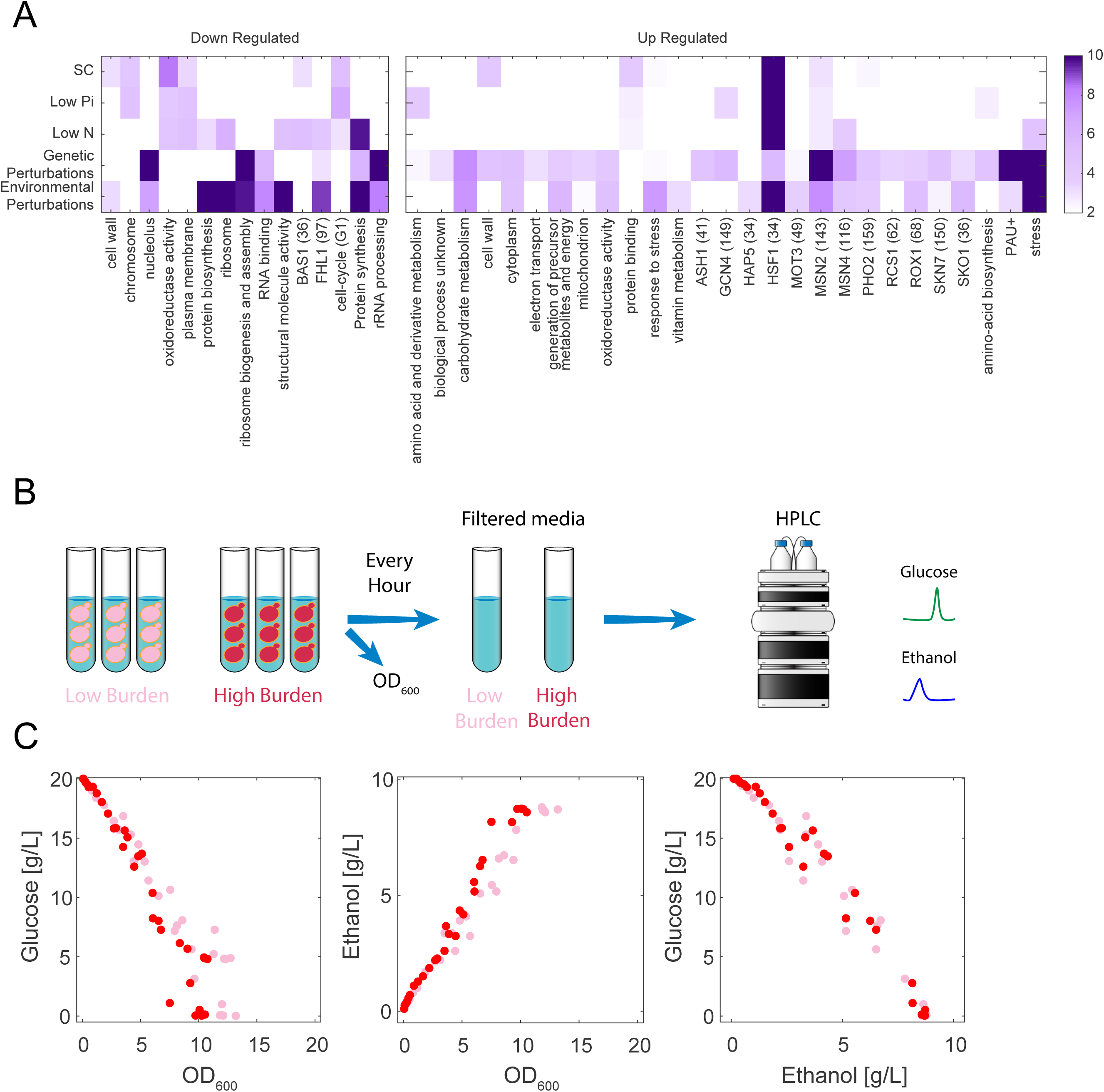
(A) GOslim, transcription factors and functional genes groups (Ihmels et al., 2002) enrichment of genes that change with the increasing burden more than ±1.5SD in each dataset. Color bar indicates log_10_(P-value). (B) HPLC experimental scheme: OD was measured and filtered media from low and high burden strains growing in SD media was measured in HPLC for Ethanol and Glucose content approximately every hour. (C) Glucose consumption and Ethanol production in low (pink) and high (red) burden strains as measured by the HPLC.

**Figure S3.**
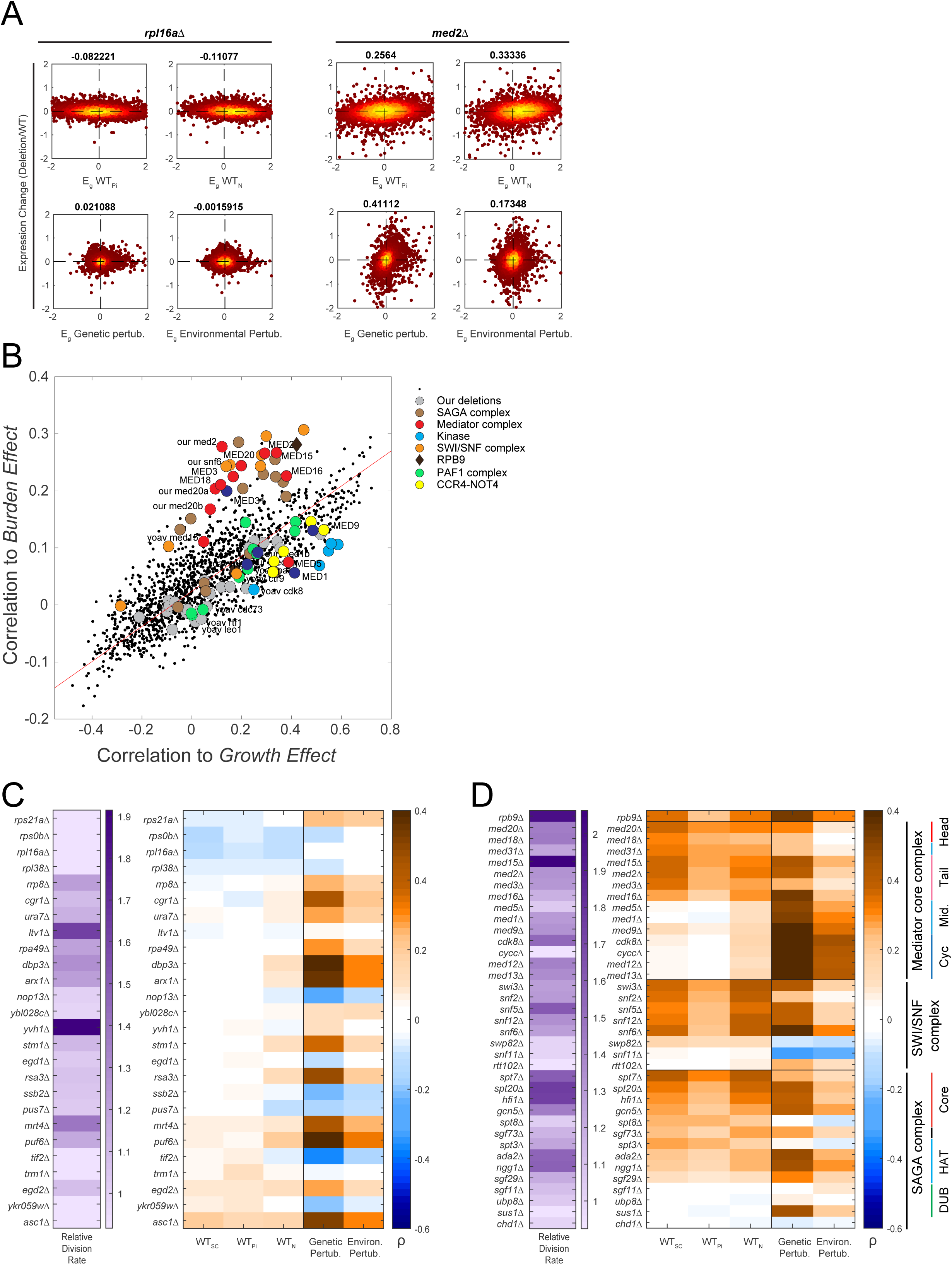
(A) The expression changes in the indicated deletion vs. the Eg in the datasets not shown in Figure 3A. (B) Correlation to Burden effect (Pearson correlation of six burden conditions averaged) is plotted vs. the Correlation to Growth effect (Pearson correlation of the two perturbations average) for different functional groups. Deletions marked with dashed circles denote verification deletions from this study and a previous study in our lab (Voichek et al., 2018). (C) Left – Reported relative growth rate to WT. Right - Ribosomal associated gene deletions correlations. (D) Right – Transcriptional initiation associated gene deletions correlations, Left – Reported relative growth rate to WT.

**Figure S4.**
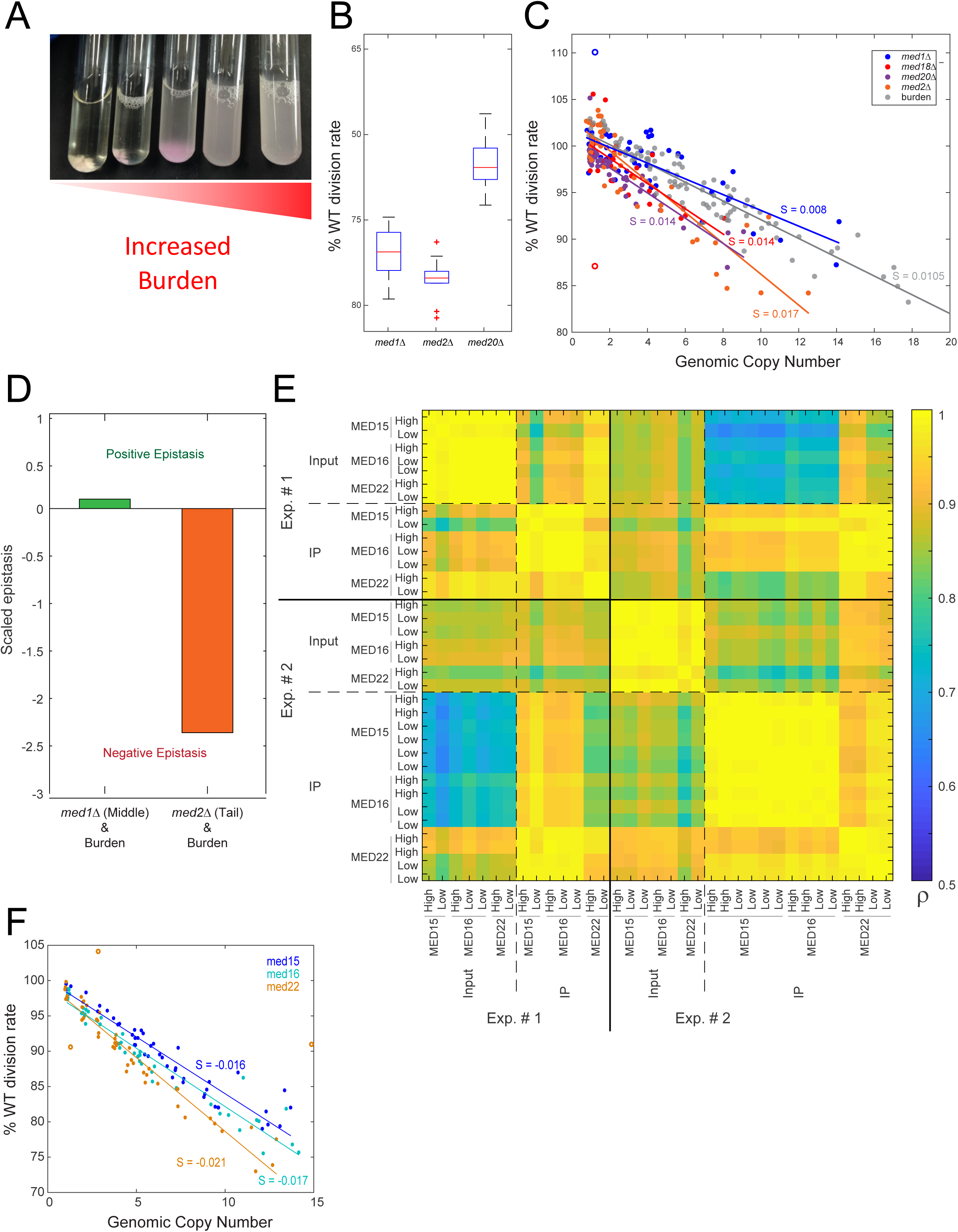
Similar effects of the burden and the Mediator phenotypes. (A) Increasing burden on the background of *cdk8*Δ. As the burden increases, flocculation decreases. (B) Relative fitness of different deletion mutants as measured in a competition assays. (C) Competition assay for Mediator deletion strains used for Figure 4C. Lines are linear fits, *S* denotes the slope of the fit. (D) Additional epistasis interaction analysis for a second competition experiment, same analysis as Figure 4C. (E) Competition assay for Mediator MYC tagged strains used for Figure 4D-F. Lines are linear fits, *S* denote the slope of the fit. MED22 was normalized such that the linear fit crossed at y=0 (F) Pearson correlation matrix of MYC-tagged Mediator subunit ChIPseq experiments.

**Figure S5.**
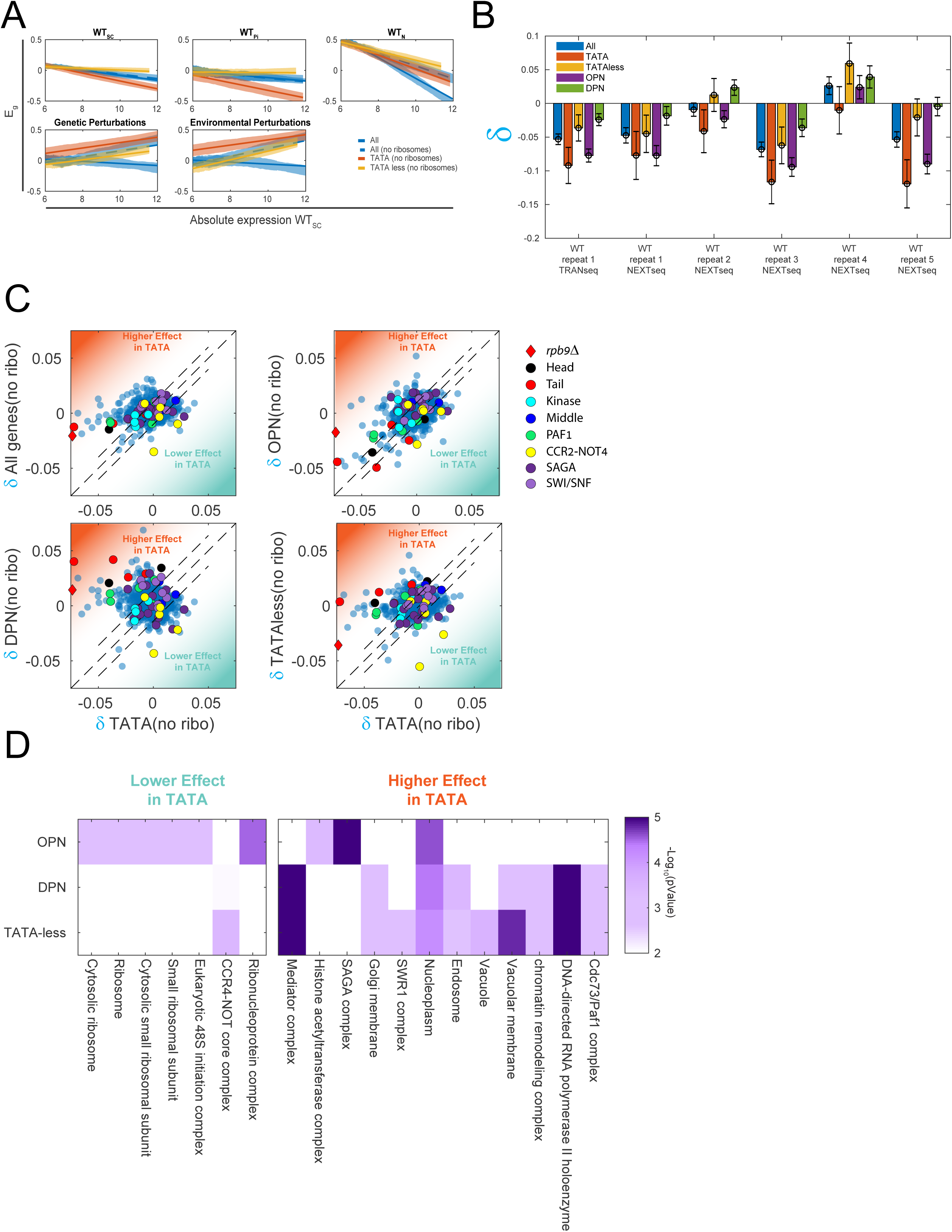
(A) TATA and OPN genes are differentially affected more than TATA-less and DPN genes in the burdened cells, but not in the deletion/environmental conditions datasets. Lines indicate a linear fit of data, shading indicates SE around the relevant fit (B) δ slope values and SE in all WT datasets shows that TATA containing genes have stronger slopes than TATA-less containing genes. (C) Comparing TATA containing gene slopes and TATA-less genes in the Genetic perturbation dataset. Each dot represents one deletion strain. (D) GO enrichment of deletion strains that have ratio of TATA<TATA-less (>1.5SD) and vice versa (box colors corresponding to C). TATA-less and DPN genes generally cluster together.

## Material and Methods

### Media and Strains

All strains of *S. cerevisiae* used in this study were constructed on the genetic backgrounds of: BY4741 (MATa his3-Δ1 leu2-Δ0 met15-Δ0 ura3-Δ0), BY4742 (MATα; his3-Δ1 leu2-Δ0 met15-Δ0 ura3-Δ0), or Y8205 (MATα; his3Δ1; LEU2Δ0; ura3Δ0; can1Δ::STE2pr-SP_his5; lyp1Δ::STE3pr-LEU2)(Brachmann et al., 1998; Tong and Boone, 2007) using standard genetic manipulations (see Table S1). Strains were grown in SC medium(Sherman and Miner, 2002) or in SC medium depleted of a specific nutrient. SC limiting media were prepared from YNB without the relevant nutrient (Low Phosphate medium - ForMedium, CYN0804, Low Nitrogen medium - BD 3101130). Phosphate depleted medium was prepared by adding phosphate in the form of KH_2_PO_4_ to a final concentration of 0.2mM. The level of potassium was preserved by adding KCl (instead of KH_2_PO_4_) in corresponding amounts. Nitrogen limiting medium was prepared from YNB without amino acids and ammonium sulfate (BD 3101130) by supplementing 50μM of ammonium sulfate and the essential amino acids. The pH values of the various media were: SC = 5.0 (except for low N, were the natural pH was about 4.9).

Deletion and double deletion strains created for validation experiments were derived from BY4741 using the LiAc/SS DNA/PEG method described(Gietz and Woods, 2002). In each strain, the gene deleted was replaced with the kanMX cassette (*geneΔ::KANMX*) using UPTAG and DNTAG primers as described in the Yeast Deletion Project (http://www.sequence.stanford.edu/group/yeast_deletion_project/usites.html). The deletion was then validated with primers A, B and kanB.

### Plasmids

p34_TDH3 and p69_TDH3 were crated as described in (Kafri et al., 2016) available at AddgeneID 127153.

### Protein burden libraries creation

Protein burden libraries were generated as described in (Kafri et al., 2016). Briefly, the pTDH3-driven mCherry plasmid was integrated into the yeast genome after linearization by the restriction enzyme MfeI. Following selection, single colonies were handpicked to create several hundred candidates. The candidates’ fluorescence levels were measured by flow cytometry. A representative library of the different fluorescence levels (indicating different copies of burden plasmid integration) was then created (each library typically contains tens of strains). Nine copies of the Myc epitope were integrated to the C terminus of Med15,16 and 22 for generation of the strains used for the ChIP analyses (plasmid pYM21 (Janke et al., 2004))

**Table S1.**
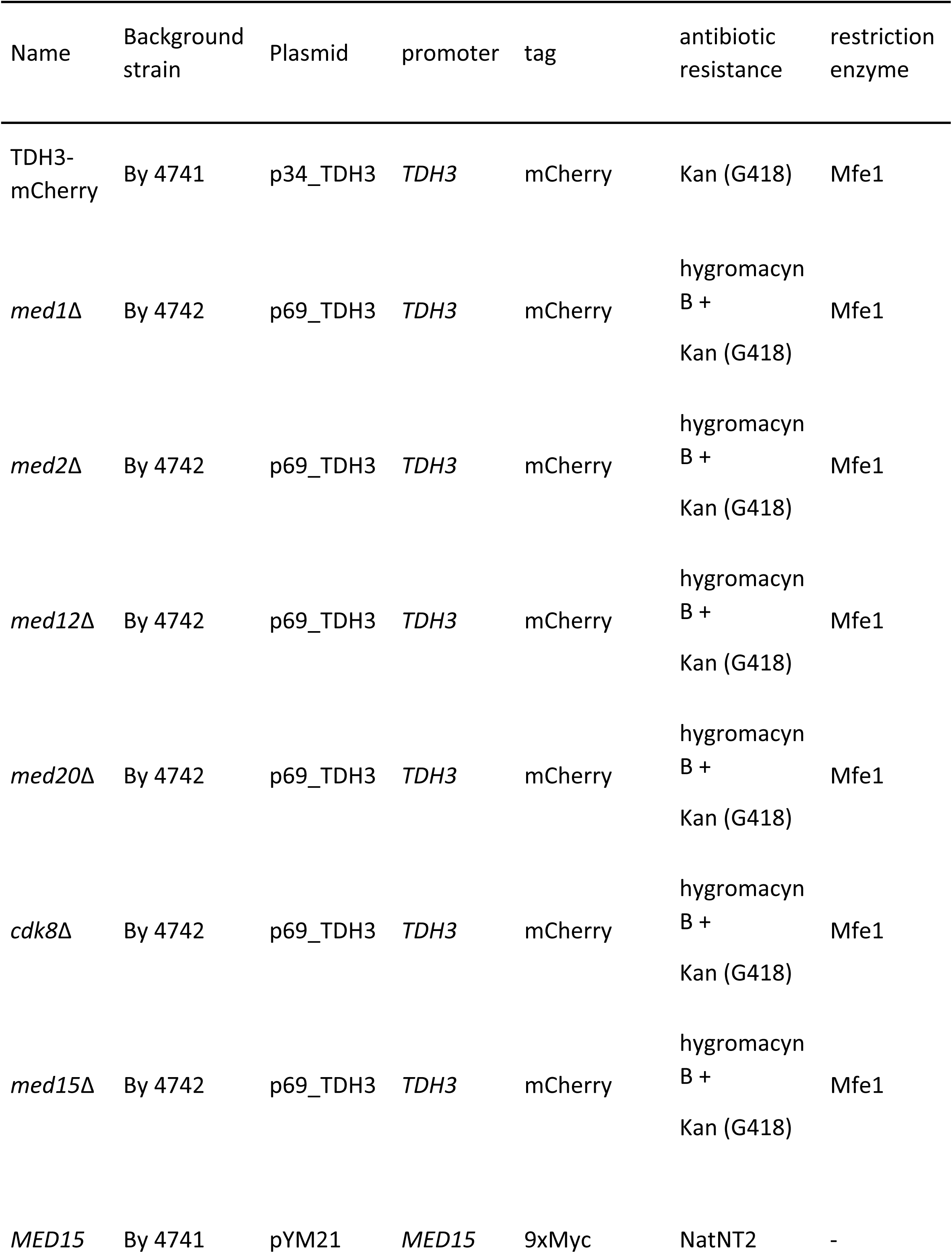
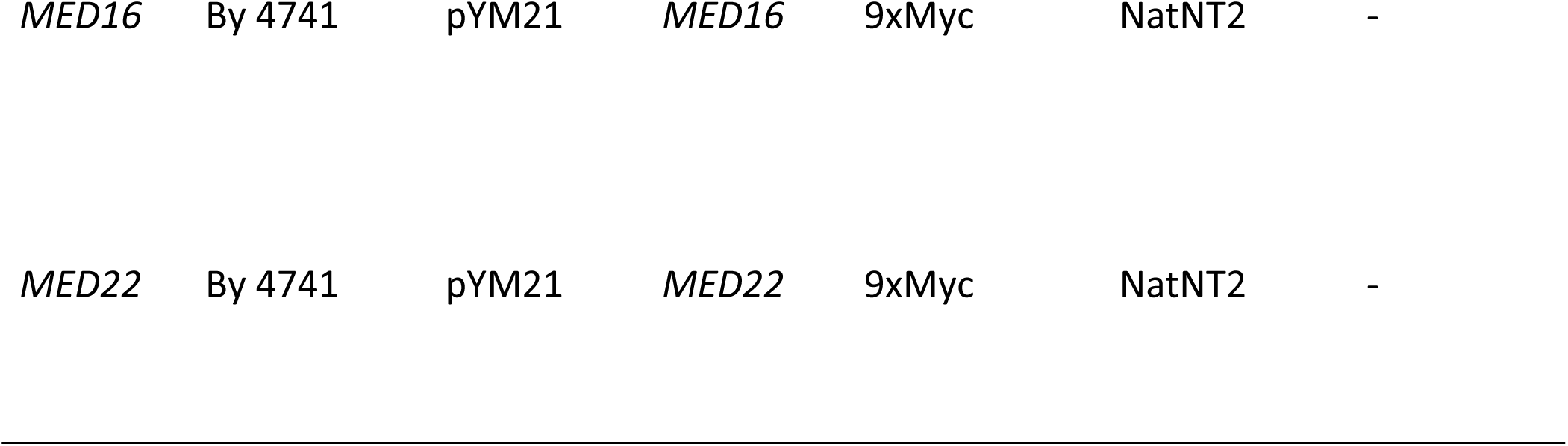
Protein burden libraries creation

### Flow cytometry

Flow cytometer measurements and analysis were done using BD LSRII system (BD Biosciences). mCherry flow cytometry was conducted with excitation at 488nm and emission at 525±25nm for GFP samples. For mCherry markers, excitation was conducted at 594nm and emission at 610±10nm. The average number of cells analyzed was 30,000.

### Competition assays

Cells were grown overnight to stationary phase. A wildtype reference GFP positive strain was then co-incubated with each of the mCherry burden strains at 30°C. The initial OD was set to ∼0.05, and the WT initial frequency was ∼50% from the total population. Following growth in the specific condition, the number of generations was calculated from the dilution factor. Frequencies of GFP versus mCherry cells were measured by flow cytometry. The cells were diluted once a day and may have reached stationary phase. A linear fit of the log_2_ for the WT frequency dynamics was used to calculate the slope for each competition assay. The relative fitness advantage is calculated from the slope divided by log2. The ‘% of WT division rate (*μ*)’ is 1 + *fitness advantage*. Each strain percentage of *μ*-*WT* was presented against its mCherry levels from the second day of the experiment or against its copy number calculated from the mCherry levels. Experiments were performed in 96 well plates.

### Epistatic calculations

#### Epistatic interactions calculations

Epistatic interactions were performed as described previously in (Segrè et al., 2005). Briefly, we calculated the scaled epistasis between the deletion mutants relative growth rate (Figure S4B) and the burden effect per one integrated copy (1 - slope of the linear fits (*S*), Figure S4C) according to the equation:

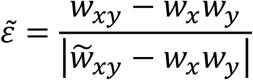

Where W_x_, W_y_, W_xy_ are the burden relative growth rate, deletion mutant relative growth rate and burden relative growth rate on the background of the deletion mutant, respectively.

Due to the burden small effects, we calculated 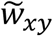 as min(*w*_*x*_, *w*_*y*_) *for w*_*xy*_ > *w*_*x*_*w*_*y*_.

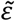 denote the epistatic interaction: 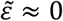, when there is no epistasis, 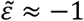 for negative epistasis and 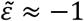 for positive epistasis.

### RNAseq transcription protocol and analysis

As described in(Voichek et al., 2016), briefly: Cells were grown to OD_600_ of 0.2-0.4 after >6hr in exponential growth and flash frozen in liquid nitrogen after centrifugation and media removal. RNA was extracted using the Nucleospin 96 RNA kit with modifications for working with yeast. Lysis was performed by mixing the cells with 300 μl lysis buffer [1M sorbitol (Sigma S1876), 100 mM EDTA 0.5 M, and 100 U/ml lyticase]. The lysis mixture was transferred to a 96-well plate that was incubated at 30° for 30 min. The plate was then centrifuged for 10 min at 3000 rpm, and the supernatant was transferred to a 96-well plate provided by the Nucleospin 96 RNA kit, followed by extraction as described in the kit protocol. Labeled cDNA was created from RNA extracts, and cDNA was barcoded and then sequenced in the Illumina HiSequation 2500 system, using a Truseq SR Cluster Kit v3-cBot-HS cluster kit and a Truseq SBS Kit v3-HS run kit (50 cycles).

### Processing and analysis of sequenced RNA

Processing and analysis of sequenced RNA was as described in Voichek *et al.* (2016). The analysis was based on the median of 6-8 exponentially growing biological repeats (SC/Low N – 8; Low Pi – 6).

To see the increase in mRNA abundance of burdened cells effect by gene abundance (Figure 5E-F), we plot (density plot) the E_g_s vs. the WT (low burden) transcription abundant. Then in (E) (Blue line) Cubic smoothing spline of data between ±1.5SD, was done and the blue shade sow – SE around the line. In (F) the slope δ was calculated using the linear fit’s slope, with the shade indicating SE.

### Flocculation assay

Flocculation assays were performed on the background of *med12*Δ as follows: several double deletion strains were created as describe above in *Strains* in addition to burden library generated as described above in *Protein burden libraries creation.* Strains were grown overnight at 30° with shaking until saturation. Next, at time point 0 the tubes were strongly vortexed for 30sec following OD_600_ measurement every few seconds as indicated in Figure S4B. OD values were normalized to time point 0.

### ChIPseq

Cells were grown overnight at 30°C with shaking to ≈OD0.6-0.8. Next, cells were washed in ice-cold PBS without Ca^++^ and Mg^++^ followed by resuspension in 2mM DSG (TS-20593, Rhenium, 50mg DSG in DMSO, PBS without Ca^++^ and Mg^++^) and agitation for 30min, RT. 1% formaldehyde was added and cells were crosslinked for an additional 5 minutes. The crosslinking was stopped by adding glycine to a final concentration of 125mM and incubating at room temperature for 5 min. Cells were then washed twice with ice-cold DDW (3800 rpm, 4^C^, 2-5 min) and flash frozen. ChIP was performed as in (Voichek et al., 2018) using Dynabeads Protein G (Invitrogen) that were incubated overnight with Myc 9E10 antibody. Cells were resuspended in lysis buffer (50mM HEPESKOH pH = 7.5, 140mM NaCl, 1mM EDTA, 1% Triton X-100, 0.1% sodium deoxycholate with freshly added Protease Inhibitor Cocktail IV (Calbiochem)) on ice and lysed mechanically with zirconium oxide beads in a BBX24-Bullet Blender (Next Advance). Lysates were then sonicated using a Diagenode Bioruptor Plus (35 cycles, high intensity, 30’’ on, 30’’ off). 30ul out of total of 600ul was taken for Input samples from each lysate. The sonicates were pre-cleared by incubation with Dynabeads Protein G incubated in binding/blocking buffer (PBSx1, 0.5% Tween, 0.5% BSA) for 1 hour at 4^c^ and subsequently incubated with antibody-coupled beads overnight. Later, lysates were washed on magnet with five rounds on lysis buffer, twice with cold buffer W1 (50 mM HEPES-KOH pH=7.5, 500 mM NaCl, 1 mM EDTA, 1% Triton X-100, 0.1% sodium deoxycholate), twice with cold buffer W2 (10 mM Tris-HCl pH=8.0, 250 mM LiCl, 0.5% NP-40, 0.5% sodium deoxycholate, 1 mM EDTA) and twice with cold TE (10 mM Tris-HCl pH=8.0, 1 mM EDTA). Then lysates were eluted with direct elution buffer (10 mM Tris-HCl pH=8.0, 1 mM EDTA, 1% SDS, 150 mM NaCl, 5 mM DTT) at 65°^C^ O.N with maximal shaking. Finally, DNA was purified by addition of 2ul RNaseA (10mg/ml), 37°C 1hr. Followed by addition of 1 ul glycogen and 2.5ul Proteinase K (20mg/ml) to each sample, 37°C 2 hr. Proteinase K Inactivation by incubation at 80°C for 20min.

### ChIP libraries

DNA from the previous step subjected to SPRI cleanup with SPRI beads 2.3x and eluted with 10mM Tris-HCL pH8. DNA libraries for Illumina NextSeq 2500 sequencing were prepared as in Yaakov et al. (2017).

### Processing and analysis of ChIP-seq

Initial processing of ChIP-seq and genomic DNA sequencing was carried out as follows: reads were aligned to a joined genome of *S. cerevisiae* (SGD, R64-1-1) and pBS69 plasmid. Genomic tracks were created from the sequence reads, representing the enrichment on each position of the joined genome. Physical fragment length was estimated by the shift best aligning the mapped sequenced reads from both ± strands, and single-end sequence reads were then lengthened accordingly (in the range of ≈100-130bp). The signal from each sample was then normalized to obtain the same total signal. Data was log_2_ transformed. Percentage occupancy on our plasmid (corresponding to amount of Mediator associated with the burden) was measured as sum of reads on the entire plasmid sequence from the total amount of reads in each sample.

### Total mRNA

*S. cerevisiae* strains and wild-type *S. paradoxus* were grown overnight at 30°C to OD_600_ ∼0.3. Cell size and count for each sample were individually assayed: Next the cultures were diluted 1:40 with 0.5M NaCl and immediately measured in Multisizer™4 COULTER COUNTER^®^ (Beckman Coulter) (Coulter, 1953). A fixed amount of ODs of *S. paradoxus* cells was added to twice as many ODs of each *S. cerevisiae* sample, such that the OD ratio between them is constant throughout the samples. The mixed samples were then flash frozen.

RNA extraction and library preparation were performed as described above and the fastq files was then processes by a pipeline for RNAseq data was created by Gil Hornung (INCPM, Weizmann Institute of Science, Israel), as described in (Herbst et al., 2017). Total reads were normalized to the ratio between the *S. cerevisiae* and *S. paradoxus* sum-of-reads and then to number of cells as measured in the experiment as described earlier.

